# Density-independent processes decouple component and ensemble density feedbacks

**DOI:** 10.1101/2021.09.19.460939

**Authors:** Corey J. A. Bradshaw, Salvador Herrando-Pérez

## Abstract

Analysis of long-term trends in abundance provide insights into population dynamics. Population growth rates are the emergent interplay of fertility, survival, and dispersal, but the density feedbacks on some vital rates (component) can be decoupled from density feedback on population growth rates (ensemble). However, the mechanisms responsible for this decoupling are poorly understood. We simulated component density feedbacks on survival in age-structured populations of long-living vertebrates and quantified how imposed nonstationarity (density-independent mortality and variation in carrying-capacity) modified the ensemble feedback signal estimated from logistic-growth models to the simulated abundance time series. The statistical detection of ensemble density feedback was largely unaffected by density-independent processes, but catastrophic and proportional mortality eroded the effect of density-dependent survival on ensemble-feedback strength more strongly than variation in carrying capacity. Thus, phenomenological models offer a robust approach to capture density feedbacks from nonstationary census data when density-independent mortality is low.

## INTRODUCTION

Compensatory density feedback describes a population’s ability to return to the environment’s carrying capacity in response to an increase in population size^1^. This phenomenon is driven by adjustments to individual fitness imposed by variation in per-capita resource availability, and associated processes of predation, competition, parasitism, and dispersal^2–5^. As survival and fertility rates ebb and flow in response to variation in population density, it is theoretically possible to detect the density-feedback signal in time series of abundance monitored at regular intervals over a sufficient period^2,6^. There is now considerable evidence that survival and fertility track population trends in many vertebrate^7–15^ and invertebrate^16–22^ species. Therefore, given the irreplaceable importance of long-term monitoring of population size in applied ecology and conservation^2^, assessing the presence of compensatory signals in censuses of population abundance remains an essential tool in the ecologist’s toolbox^23^.

The family of self-limiting population-growth models including logistic growth curves (‘phenomenological models’ hereafter) are convenient for describing density-feedback signals in abundance time series^3^. These models use census data to quantify the net effect of population size *N* on the rate of instantaneous growth (*r*)^24^. Expressed as a proportional change in *N* between two time (*t*) steps (e.g., years or generations), the assumption is that *r* = log*_e_*(*N*_*t*+1_/*N_t_*) summarises the combination or ‘ensemble’^2^ of all feedback mechanisms operating on individual ‘component’ demographic rates^25^. The problem is that population growth rates can be insensitive to variation in particular demographic rates^26–28^. Thus, across 109 observed censuses of bird and mammal populations, the strength of ‘component density feedback’ (on demographic rates) explained only < 10% of the strength of ‘ensemble density feedback’ (on population grow rate) using phenomenological models and after controlling for time-series length and body size^2^. The reasons for such decoupling are not well understood.

Determining the partial effects of different underlying mechanisms responsible for the decoupling of component and ensemble density feedbacks is most often impossible for real abundance time series. This analytical limitation occurs because the multiple, density-dependent and -independent mechanisms generating population fluctuations change themselves through time — a condition known as ‘nonstationarity’^29^. We therefore constructed stochastic, age-structured populations with known, component density feedback on survival and imposed nonstationarity to population size via multiple demographic scenarios emulating density-independent mortality and variation in carrying capacity through time. We then simulated multiannual time series of abundance from those populations and estimated the strength of ensemble density feedbacks from these. Our prediction was that ensemble density feedbacks should track component feedbacks if survival has a demographic impact, mediated by population size, on the population growth rate of long-lived vertebrates, while our demographic framework allowed the quantification of true and false detection of ensemble density feedbacks.

## METHODS

Our overarching aim was to simulate populations of long-living species and their time series of abundance with various sources of nonstationarity. We describe below the set of test species, the simulation of the base population model, component density feedbacks on survival and time series of population abundance, the demographic scenarios considered, and the phenomenological models used to quantify ensemble density feedbacks.

### Test species

As the variability in population growth rates is driven primarily by survival rates for slower life-history species of mammals^30,31^ and birds^32^, we parameterised the simulated populations to characterise the plausible dynamics of 21 long-lived species of extant (*n* = 8) and extinct (*n* = 13) Australian vertebrates from five taxonomic/functional groups (herbivore vombatiformes and macropodiformes, large omnivore birds, carnivores, and invertivore monotremes), spanning mean adult body masses of 1.7–2786 kg and generation lengths of 2.3–21 years^33^ (Table 1). These species differ in their resilience to environmental change, and represent the slow end of the slow-fast continuum of life histories^34^ where high survival rates make it possible that reproductive efforts are dispersed over the lifetime of individuals^35^. A full justification of the selection of our test species can be found in reference^33^.

**TABLE 1.** Taxonomy and life-history characteristics of the 21 test species (all native to Australia) used to simulate age-structured populations and time series of population abundance. abb = abbreviation of scientific name, *M* = body mass (kg), *GL* = generation length (years), *q* = projection length (years)^33^.

| taxonomic/functional group | species | abb | $M$ | $GL$ | $q$ | status |
| --- | --- | --- | --- | --- | --- | --- |
| herbivore<br>vombatiformes | *Diprotodon optatum* | DP | 2786 | 18.1 | 724 | extinct |
|  | *Palorchestes azael* | PA | 1000 | 15.1 | 604 | extinct |
|  | *Zygomaturus trilobus* | ZT | 500 | 13.2 | 528 | extinct |
|  | *Phascolonus gigas* | PH | 200 | 10.7 | 428 | extinct |
|  | *Vombatus ursinus* | VU | 25 | 10.0 | 400 | extant |
| herbivore<br>macropodiformes | *Procoptodon goliah* | PG | 250 | 8.3 | 332 | extinct |
|  | *Sthenurus stirlingi* | SS | 150 | 8.1 | 324 | extinct |
|  | *Protemnodon anak* | PT | 130 | 7.8 | 312 | extinct |
|  | *Simosthenurus occidentalis* | SO | 120 | 7.8 | 312 | extinct |
|  | *Metasthenurus newtonae* | MN | 55 | 6.0 | 240 | extinct |
|  | *Osphranter rufus* | OR | 25 | 5.5 | 220 | extant |
|  | *Notamacropus rufogriseus* | NR | 14 | 6.3 | 252 | extant |
| large omnivore birds | *Genyornis newtoni* | GN | 200 | 20.0 | 800 | extinct |
|  | *Dromaius novaehollandiae* | DN | 55 | 5.9 | 236 | extant |
|  | *Alectura lathamii* | AL | 2.2 | 6.8 | 272 | extant |
| carnivores | *Thylacoleo carnifex* | TC | 110 | 9.1 | 364 | extinct |
|  | *Thylacinus cynocephalus* | TH | 20 | 5.2 | 208 | extinct |
|  | *Sarcophilus harrisii* | SH | 6.1 | 3.1 | 124 | extant* |
|  | *Dasyurus maculatus* | DM | 2 | 2.3 | 92 | extant |
| invertivore<br>monotremes | *Megalibgwilia ramsayi* | MR | 11 | 16.4 | 656 | extant |
|  | *Tachyglossus aculeatus* | TA | 4 | 14.1 | 564 | extant |

\* extant in Tasmania, currently extinct in mainland Australia

### Base (age-structured) population model

The population model for each test species was a stochastic (parameters resampled within their uncertainty bounds) Leslie transition matrix (**M**) following a pre-breeding design, with *ω*+1 (*i*) × *ω*+1 (*j*) elements (representing ages from 0 to *ω* years) for females only, where *ω* represents maximum longevity. Fertility (*m_x_*) occupied the first row of the matrix, survival probabilities (*S_x_*) occupied the sub-diagonal, and the final diagonal transition probability (**M***_i,j_*) was *S_ω_* for all species — except *Vombatus ursinus* (VU; common wombat), *Thylacinus cynocephalus* (TC; thylacine) and *Sarcophilus harrisii* (SH; devil) for which we set *S_ω_* = 0 to limit unrealistically high proportions of old individuals in the population given the evidence for catastrophic mortality at *ω* for the latter two species^36–38^. Multiplying **M** by a population vector **n** estimates total population size (Σ**n**) at each forecasted time step^39^. The base model was parameterised with **n**_0_ =*AD***Mw**, where **w** is the right eigenvector of **M** (stable stage distribution), and *A* is the surface area of the study zone (*A* = 250,000 km^2^) so that the species with the lowest **n**_0_ would have an initial population of at least several thousand individuals at the start of the simulations. Based on theoretical equilibrium densities (*D*, km^-2^) calculated for each taxon^33^, the species-specific carrying capacity *K* = *DA.*

### Density feedback on survival

We simulated a compensatory density-feedback function by forcing a reduction modifier (*S*_red_) of the *S_x_* vector in each model according to Σ**n**:

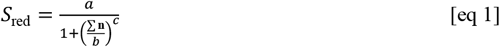

where the *a*, *b*, and *c* constants for each species are adjusted to produce a stable population on average over 40 generations (40⌊*G*⌉; see below)^40,41^. This formulation avoided exponentially increasing populations, optimised transition matrices to produce parameter values as close as possible to the maximum potential rates of instantaneous increase (*r*_m_)^33^, and so ensured that long-term population dynamics were approximately stable at the species-specific *K* (see previous section). Here,

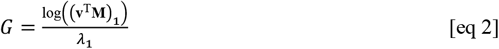

(**v**^T^**M**)_1_ is the dominant eigenvalue of the reproductive matrix **R** derived from **M**, and **v** is the left eigenvector^39^ of **M**. Thus, the total projection length in years (*q*) varied across the 21 test species, from 92 (*Dasyurus maculatus;* DM; spot-tailed quoll) to 800 (*Genyornis newtoni;* GN; mihirung) years (median = 324 years with 95 % interquartiles of [108, 762] years; Table 1), with one value of abundance being simulated per year. In each iteration and annual time step, the *S_x_* vector was *β*-resampled assuming a 5% standard deviation of each *S_x_* and a Gaussian-resampled *m_x_* vector. We deliberately avoided applying density-feedback functions to fertility to isolate the component feedback to a single demographic rate.

### Nonstationarity

We added nonstationarity to our base population model through a catastrophic (density-independent) mortality function to account for the probability of a catastrophic event (*C*) scaling to generation length among vertebrates^42^:

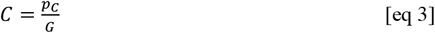

where *p_C_* = probability of catastrophe set at 0.14 given this is the mean probability per generation observed across vertebrates^42^. Once invoked at probability *C*, a *β*-resampled proportion centred on 0.5 to the *β*-resampled *S_x_* vector induced a ~ 50% mortality event for that year^43^. A catastrophic event is defined as “… any 1-yr peak-to-trough decline in estimated numbers of 50% or greater”^42^. The catastrophic function recreates the demographic effects of a density-independent process such as extreme weather events, fires, disease outbreaks, or human harvest. However, we considered the process here as a standard perturbation in all models, and then added specific types of additional perturbations per scenario (see demographic scenarios below).

### Abundance time series

From the base models (parameterised to incorporate age structure, density feedbacks on survival, and catastrophic events in the Leslie matrices as described above), we generated multiannual abundance time series up to 40 ⌊*G*⌉ for each species. We standardised projection length to 40 ⌊*G*⌉ because there is strong evidence that the length of a time series (*q*) dictates the statistical power to detect an ensemble density-feedback signal in logistic growth curves^6^. Here, we summed the **n** vector over all age classes to produce a total population size *N_t,i_* for each year *t* of each iteration *i*. We rejected the first ⌊*G*⌉-equivalent years of each projection as a burn-in to allow the initial (deterministic) age distribution to calibrate to the stochastic expression of stability under compensatory density feedback.

To ascertain the degree of nonstationary in the simulated abundance time series, we used Turchin’s^29^ definition of nonstationarity as temporally variant mechanisms generating population fluctuations. In that conceptual context, we calculated the mean and variance of return time (*T*_R_) — defined as the time required to return to equilibrium following a disturbance^44^ — for each abundance time series as:

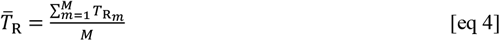

where 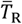 is the mean *T*_R_ across *M* steps of the time series. For each *m*^th^ time step,

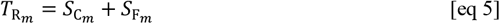

where: *S*_C*_m_*_ is the number of complete time steps taken before reaching *T*_R*_m_*_, and *S*_F*_m_*_ is the fraction of time required to reach *T*_R*_m_*_ in the *M*^th^ (final) step:

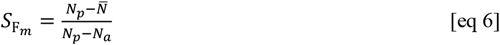

where 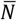 is the mean of the abundance time series (a proxy for *K*), *N_p_* is the population size prior to crossing 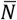, and *N_a_* is the population size after crossing 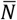.

The variance of *T*_R_ is:

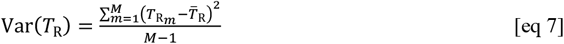

Thus, when 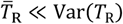 (i.e., 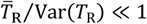), the time series is considered to be highly nonstationary^44^.

### Demographic scenarios

We generated 10,000 abundance time series over 40 ⌊*G*⌉ for each test species in each of nine demographic scenarios that combined different types and magnitudes of nonstationarity in the form of density-independent (catastrophic and proportional) mortality and variation in carrying capacity (*K*) through time. Each times series represented the idiosyncratic demography of a unique population occupying an area of 250,000 km^2^ with zero dispersal (see above).

We split the nine scenarios into two main groups: (**1**) eight testing the probability of a false negative (reduced detection of ensemble density feedback when a component feedback on survival existed), and (**2**) one testing the probability of a false positive (evidence of ensemble density feedback when a component feedback on survival was absent) (see details in Table 2). The false-negative scenarios included three subcategories: (**1.1**) ***i***. fixed *K* with no perturbations other than the stochasticity imposed by resampling demographic rates in the Leslie matrices; (**1.2**) fixed *K* with generationally scaled catastrophes centred on 50% mortality ***ii***. leading to 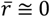, ***iii.*** as in *ii*, but with an additional, single ‘pulse’ perturbation of 90% mortality applied across the entire age structure at 20 generations, ***iv.*** a ‘harvest’-like process where a consistent proportion of individuals is removed from the **n** vector at each time step to produce 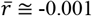 (i.e., weak, monotonic decline in average population size), or ***v.*** as in *iv*, but where the resultant 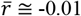 (i.e., strong, monotonic decline in average population size); and (**1.3**) *K* fluctuations with ***vi.*** stochastically resampled *K* with a constant 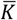 and a constant variance (via resampling the *b* parameter in equation [1]), ***vii.*** as in *vi*, but where the resampling variance doubles over the projection interval (via a linear increase in the standard error used to resample the *b* parameter in equation [1]), and ***viii***. as in *vi*, but where *K* declines at a rate of 0.001 over the projection interval (via decreasing the *b* parameter in equation [1]). **2**. The false-positive scenario **2*ix.*** tested for false positives in the ensemble signal by imposing a density-independent mortality via an increase in the probability of catastrophe *p_C_* in equation [3] to produce 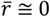 over 40⌊*G*⌉ In that scenario, we removed the component density-feedback on survival (i.e., setting *S*_red_ = 1) — theoretically, populations lack a carrying capacity in the absence of density feedbacks.

**TABLE 2.** Demographic scenarios to quantify the detection of ensemble density-feedback signals in time series of abundance using phenomenological models (logistic growth curves) if a component density feedback on survival is present (1. H_0_: false negatives), or absent (2. H_0_: false positives). All scenarios were simulated over 40 generations across 21 vertebrate species. Time series obtained from simulated age-structured populations (Leslie matrices) occupying 250,000 km^2^ with no dispersal. *G* = generation, *N* = population abundance, *K* = carrying capacity; 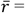 long-term mean instantaneous rate of population change, SD = standard deviation. See test species in Table 1.

| scenario | catastrophe type | description |
| --- | --- | --- |
| **1. $H_0$: false negatives (component feedback)** |  |  |
| *1.1 no catastrophic mortality or fluctuation in $K$* |  |  |
| i. $K_{\text{fixed}}, \bar{r} \cong 0$ | none | stochastically resampled survival rates in age-structured population |
| *1.2 catastrophic mortality (50%) and stable $K$* |  |  |
| ii. $K_{\text{fixed}}, \bar{r} \cong 0$; sustained catastrophic mortality | generationally scaled | as in *i*, but with catastrophes |
| iii. $K_{\text{fixed}}, \bar{r} \cong 0$; pulsed catastrophic mortality | generationally scaled | as in *ii*, but with a single 90% mortality pulse implemented at $20G$ |
| iv. $K_{\text{fixed}}, \bar{r} \cong -0.001$; sustained proportional mortality | generationally scaled | as in *ii*, but with proportional removal of individuals from the $\mathbf{n}$ vector such that $\bar{r} = -0.001$ (slowly declining population) |
| v. $K_{\text{fixed}}, \bar{r} \cong -0.01$; sustained proportional mortality | generationally scaled | as in *iv*, but where $\bar{r} = -0.01$ (rapidly declining population) |
| *1.3 catastrophic mortality (50%) and fluctuation in $K$* |  |  |
| vi. $K_{\text{stochastic}}, \bar{r} \cong 0$ | generationally scaled | as in *ii*, but normally distributed $K$ varying randomly at each time step (SD = 5%) |
| vii. $K_{\text{stochastic}}$ with increasing variance; $\bar{r} \cong 0$ | generationally scaled | as in *vi*, but variance in $K$ increased linearly from 5% to 10% |
| viii. $K_{\text{stochastic}}$ declining, forcing $\bar{r} < 0$ | generationally scaled | as in *vi*, but $K$ also decreases on average at a rate of -0.001 |
| **2. $H_0$: false positives (no component feedback)** |  |  |
| ix. no $K$; $\bar{r} \cong 0$ | temporally scaled | probability of catastrophe increased over time such that $\bar{r} \cong 0$ (~ average stability) |

### Ensemble density feedbacks

After generating 10,000 time series for each of the 21 species following the nine demographic scenarios (totalling 189,000 individual time series), we applied phenomenological models to each time series to test the statistical *evidence* for an ensemble compensatory density feedback, as well as quantify the *strength* of that feedback. Our phenomenological models included four variants of the general logistic growth curve^45^ following reference^6^:

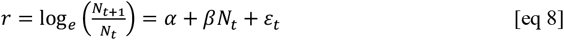

where *N_t_* = population size at time *t, α* = intercept, *β* = strength of ensemble density feedback, and *ε_t_* = Gaussian random variable with a mean of zero and a variance *σ*^2^ reflecting uncorrelated stochastic variability in the instantaneous rate of population change *r*. Our first two models are simple density-independent models (DI): (1) random walk, where *α* = *β* = 0, and (2) exponential growth, where *β* = 0. The second two variants are density-dependent or density-feedback models (DF): (3) Ricker-logistic ^46^, and (4) Gompertz-logistic^47^, where *Nt* on the right side of equation [8] is replaced with log*_e_*(*N_t_*). The latter two models represent alternative situations where population growth rate varies in response to unit (Ricker) or order-of-magnitude (Gompertz) changes in population size^1^.

After fitting each of the four phenomenological models to each time series, we calculated their relative likelihood by means of the Akaike’s information criterion (AIC) corrected for finite number of samples (AIC*_c_*). We then expressed the *evidence* for an ensemble density-feedback signal Pr(DF) as the sum of AIC*_c_* weights (*w*AIC*_c_* = model probability)^48^ for the Ricker- and Gompertz-logistic models (i.e., Σ*w*AIC*_c_*-DF), and the *evidence* for a lack of such signal as the sum of AIC*_c_* weights for random walk and exponential growth (i.e., Σ*w*AIC*_c_*-DI). This follows the logic that if *β* ≠ 0 between *r* and *N_t_* (Ricker) *or* log*_e_*(*N_t_*) (Gompertz) is more likely than *β* = 0 (random walk and exponential growth), then there is stronger statistical support for an ensemble density feedback in the time series than not (i.e., Σ*w*AIC*_c_*-DF > Σ*w*AIC*_c_*-DI implies Pr(DF) > 0.5).

We estimated the *strength* of the ensemble density-feedback signal as the negative value of 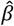 estimated from the Gompertz-logistic model. We used the Gompertz-logistic 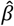, instead of the Ricker-logistic 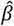, to estimate this strength because only the former characterises the multiplicative nature of demographic rates^2,49^. To compare the component density feedback applied to survival in the stochastic age-structured models to the ensemble density feedback estimated from the abundance time series under the nine demographic scenarios, we plotted the negative value of Gompertz 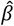 relative to 1 – *S*_red_ across all 21 species modelled.

We tested the correlation between ensemble and component density-feedback strength, and between ensemble strength and the degree of stationarity, across species by calculating a bootstrapped estimate of Spearman’s correlation *ρ* (treating relative differences in the metrics as ranks). We uniformly resampled 10,000 times from the 95% confidence interval of each metric for each species and demographic scenario, calculating *ρ* in turn, and then calculating the median and 95% confidence interval of *ρ*. The relationships between ensemble and component density-feedback strength (as well as between ensemble strength and stationarity) showed some non-linearity, so we also fit simple exponential plateau models of the form *y* = *y*_max_ - (*y*_max_ - *y*_0_)e*^-kx^* to these relationships. Here, *y*_0_ is the starting value of component strength, *y*_max_ is the maximum component strength (- Gompertz 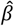), *k* = rate constant (in units of *x*^-1^), and *x* is the component strength (1 – *S*_red_).

## RESULTS

### Statistical evidence for density feedback

For each test species, when the simulated populations were subjected to a compensatory density feedback on survival (age-structured Leslie matrices), the median probability for a statistical signal of ensemble compensatory density-feedback (Pr(DF) = Σ*w*AIC*_c_*-DF; see Materials and methods) across 10,000 times series of abundance was near unity (> 0.99) for the stable 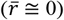 trajectories and most demographic scenarios (Fig. S1–S2 and S3 for probability density plots of Pr(DF) across scenarios and the bootstrapped mean Pr(DF) per species and scenario, respectively). Only the declining stochastic *K* scenario (1.3*viii*) had a slightly smaller median Pr(DF) at 0.95. For the false-positive scenario (2*ix*), the median Pr(DF) was 0.322. Generally, the extant dasyurid *S. harrissii* (SH; devil) and the flightless bird *Dromaius novaehollandiae* (DN; emu) had the weakest evidence for density feedback across the different scenarios (Fig. S3).

In summary, if a component density feedback was present, the phenomenological models mostly detected the ensuing ensemble feedback (true positive) — regardless of whether a simulated population was perturbed via density-independent removal of individuals, or altered *K* dynamics — in > 9 of every 10 time series; while false positives (component feedback absent, ensemble feedback detected) occurred in < 4 of every 10 times series.

### Degree of simulated stationarity

The addition of the generationally scaled 50% catastrophic (density-independent) mortality reduced stationarity from a median of 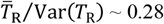 (scenario 1.1*i*) to ~ 0.08 (scenario 1.2*ii*) (Fig. 1A). The scenarios imposing a catastrophic 90% mortality as a pulse at 20 generations (1.2*iii*), or additional proportional mortality driving a moderately (1.2*iv*; 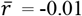) or rapidly (1.2*v*; 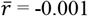) declining population over 40 generations, all reduced stationarity by approximately the same amount relative to the scenario without catastrophic mortality (1.1*i*) (Fig. 1C). For the scenarios emulating fluctuations in *K* (1.3*vi*–*viii*), adding stochasticity to *K* slightly increased stationarity relative to a fixed-*K* scenario (Fig. 1E). Only when the stochastic *K* was forced to decline (scenario 1.3*viii*), the abundance time series became highly nonstationary (Fig. 1E). The false-positive scenario (2.*ix*) resulted in negligible change to stationarity when comparing populations experiencing (Fig. 2A), or not experiencing (Fig. 2B), a component density feedback on survival.

**FIGURE 1.**
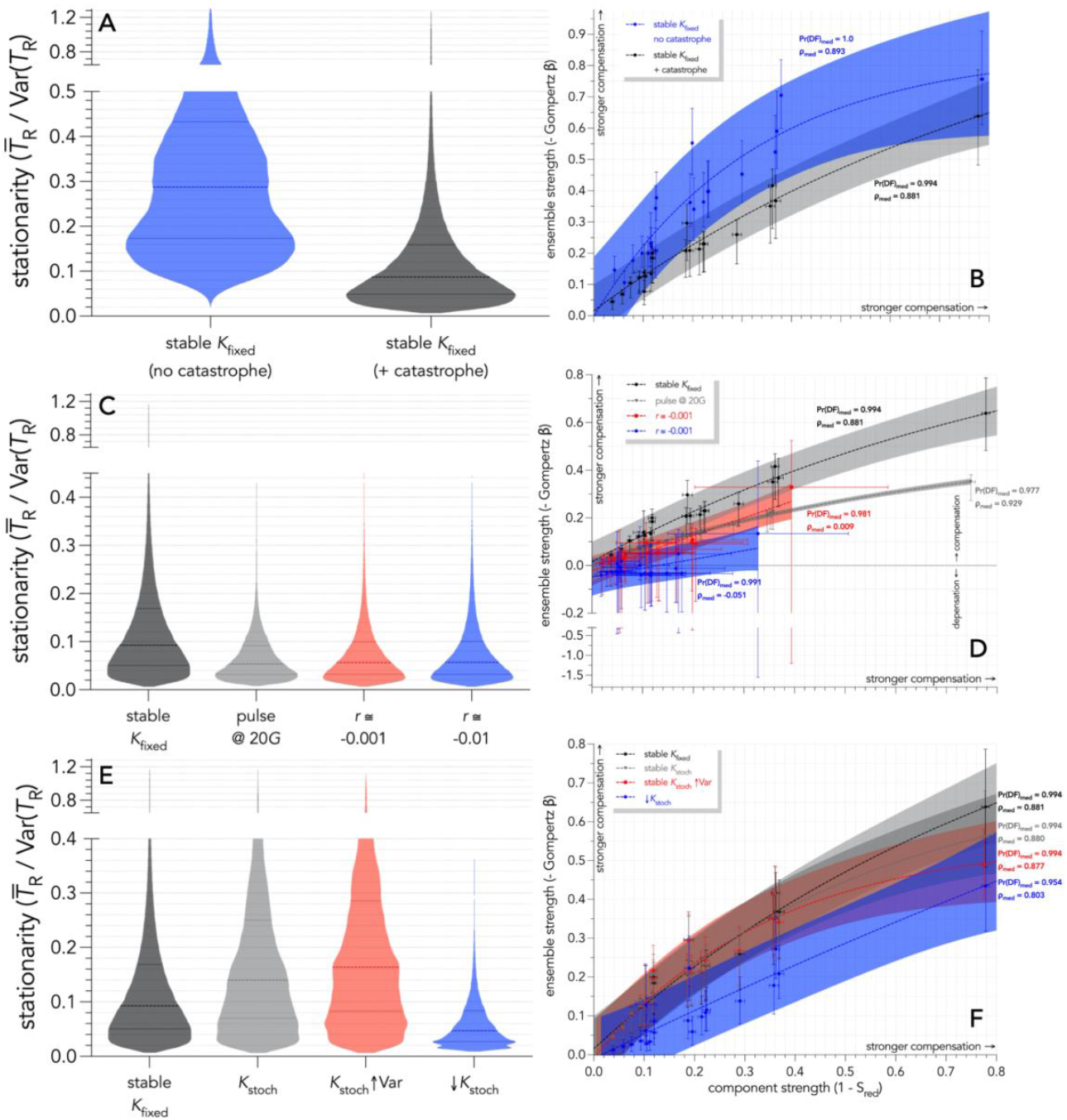
(**A**, **C**, **E**) Truncated violin plots showing the distribution of the stationarity index 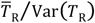 across 10,000 times series of population abundance per species and all 21 test species (see list in Table 1) obtained from age-structured populations subjected to a compensatory component density feedback on survival over 40 generations, according to nine demographic scenarios (detailed in Table 2). (**B**, **D**, **F**) Relationship between strength of ensemble (slope coefficient *β* of the Gompertz-logistic model × [−1]) and component (1 – the modifier *S*_red_ on survival) density feedback. (**A**-**B**) Scenarios without (blue: scenario 1.1*i*) and with (grey: scenario 1.2*ii*) generationally scaled 50% catastrophic (density-independent) mortality. (**C**-**D**) Stable projections with carrying capacity (*K*) fixed (darker grey; scenario 1.2*ii*), a pulse disturbance of 90% mortality at the first 20 generations (20*G*; lighter grey; scenario 1.2*iii*), weakly declining (*r* ≅ −0.001; red; scenario 1.2*iv*), and strongly declining (*r* ≅ 0.01; blue; scenario 1.2*v*). (**E**-**F**) Stable projections with *K* fixed (darker grey; scenario 1.2*ii*), varying stochastically (*K*_stoch_) around a constant mean with a constant variance (lighter grey; scenario 1.3*vi*), varying stochastically with a constant mean and an increasing variance (*K*_stoch_↑Var; red; scenario 1.3*vii*), and varying stochastically with a declining mean and a constant variance (↓*K*_stoch_; blue; scenario 1.3*viii*). The fitted curves across species are exponential plateau models of the form *y* = *y*_max_ - (*y*_max_ - *y*_0_)e*^-kx^*. Shaded regions represent the 95% prediction intervals for each type. Also shown are the mean probabilities of median density feedback (Pr(DF): sum of the Akaike’s information criterion weights for the Ricker- and Gompertz-logistic models across time series (Σ*w*AIC*_c_*-DF). Compensation implies that survival and population growth wane as population abundance rises, and 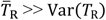 implies high stationarity.

**FIGURE 2.**
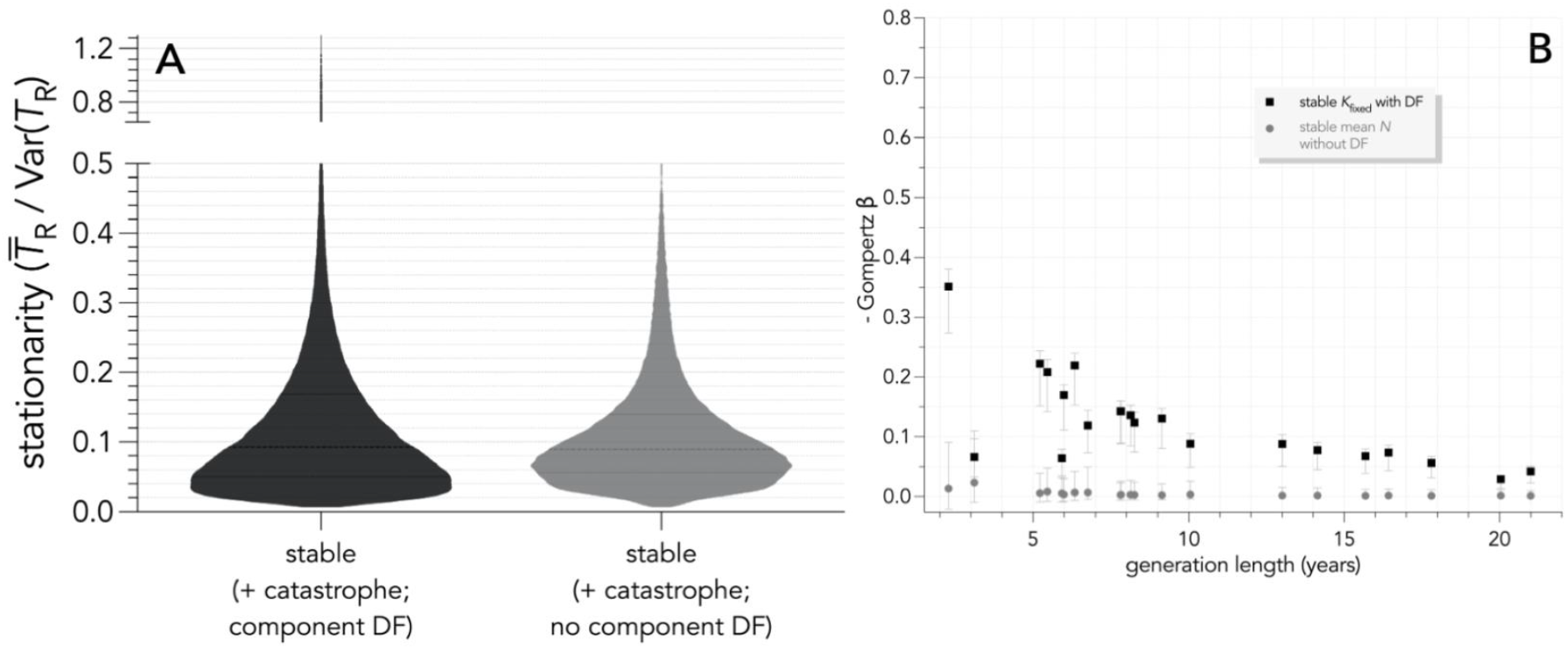
(**A**) Truncated violin plots showing the distribution of the stationarity index 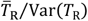 across 10,000 times series of population abundance per species and all 21 species (see species list in Table 1) obtained from age-structured populations subjected to a compensatory component density feedback on survival over 40 generations, according to two demographic scenarios (detailed in Table 2). Demographic scenarios include carrying capacity (*K*) fixed with (darker grey, scenario 1.2*ii*) and without (lighter grey, scenario 2*ix*) component compensatory density-feedback on survival, the latter including an increase in the probability of 50% catastrophic (density-independent) mortality to produce stable population growth rates around 0 (see scenarios in Table 2). (**B**) Relationship between strength of ensemble (slope coefficient *β* × [−1] of the Gompertz-logistic model) and generation length (years) across the 21 species. Probabilities of density feedback (Pr(DF) = sum of the Akaike’s information criterion weights for the Ricker and Gompertz models) calculated across simulations gave median Pr(DF) = 0.994 and 0.322 for the two stable scenarios without and with component feedback on survival, respectively.

### Strength of density feedback

While the magnitude of statistical evidence for density feedback was largely invariant across all demographic scenarios including a component density feedback on survival (Fig. S1 and S2; see above), the estimated strength of the ensemble density feedback (-Gompertz *β*, see Materials and methods) was highly sensitive to the type of perturbation the population experienced. The addition of the generationally scaled 50% catastrophic (density-independent) mortality under a fixed *K* (scenarios 1.1*i vs*. 1.2*ii*) reduced the correlation (median *ρ* = 0.893 and 0.881, respectively) and slope between ensemble feedback strength and component feedback strength (1 – *S*_red_) across the 21 test species (Fig. 1B). The catastrophic pulse scenario (1.2*iii*) returned the closest correlation (median *ρ* = 0.929) between ensemble and component feedback strengths, although it also depressed the slope of the relationship relative to the *K*_fixed_ scenario (Fig. 1D). These correlations were weakest for the 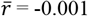 and 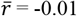 scenarios (1.2*v*–*vi*; median *ρ* = 0.009 and −0.051, respectively), which also captured a signal of depensation (population growth rate increases with population size) in some abundance time series (Fig. 1D). For the demographic scenarios emulating fluctuations in *K* (1.3), the correlation between unit change in ensemble and component density feedback strength was generally higher than those where 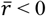 (Fig. 1F; median *ρ* ranging from 0.803 to 0.881), with the strongest mismatch occurring when *K* declined by a rate of 0.001 (scenario 1.3*viii*) (Fig. 1F; see also Fig. S4). For the false-positive scenario (2*ix*), all estimated ensemble feedback strengths enveloped 0 (Fig. 2B), meaning that the estimated slopes of the *r* ~ log*_e_*(*N_t_*) relationships could not be differentiated from zero.

Overall, when an ensemble density feedback was detected from time series of abundance, density-independent mortality eroded the extent by which true compensatory density feedbacks on survival translated into an ensemble compensatory density feedback in population trends more than fluctuations in *K*, with the most faulty outcome in fact inferring depensatory population growth rates from some populations only experiencing density compensation on survival.

On the other hand, the stationarity metric 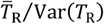 was a weak (median *ρ* = 0.547, −0.086, and −0.113 for the pulse, 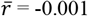, and 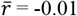 scenarios, respectively) predictor of the estimated strength of ensemble feedback when density-independent mortality was imposed (Fig. 3). However, stationarity was a reasonable (median *ρ* = 0.756, 0.786, and 0.844 for the *K*_stochastic_, *K*_stochastic_ with increasing variance, and declining *K*_stochastic_ scenarios, respectively) predictor of the ensemble signal for the fluctuating *K* scenarios (Fig. 4; see also Fig. S4).

**FIGURE 3.**
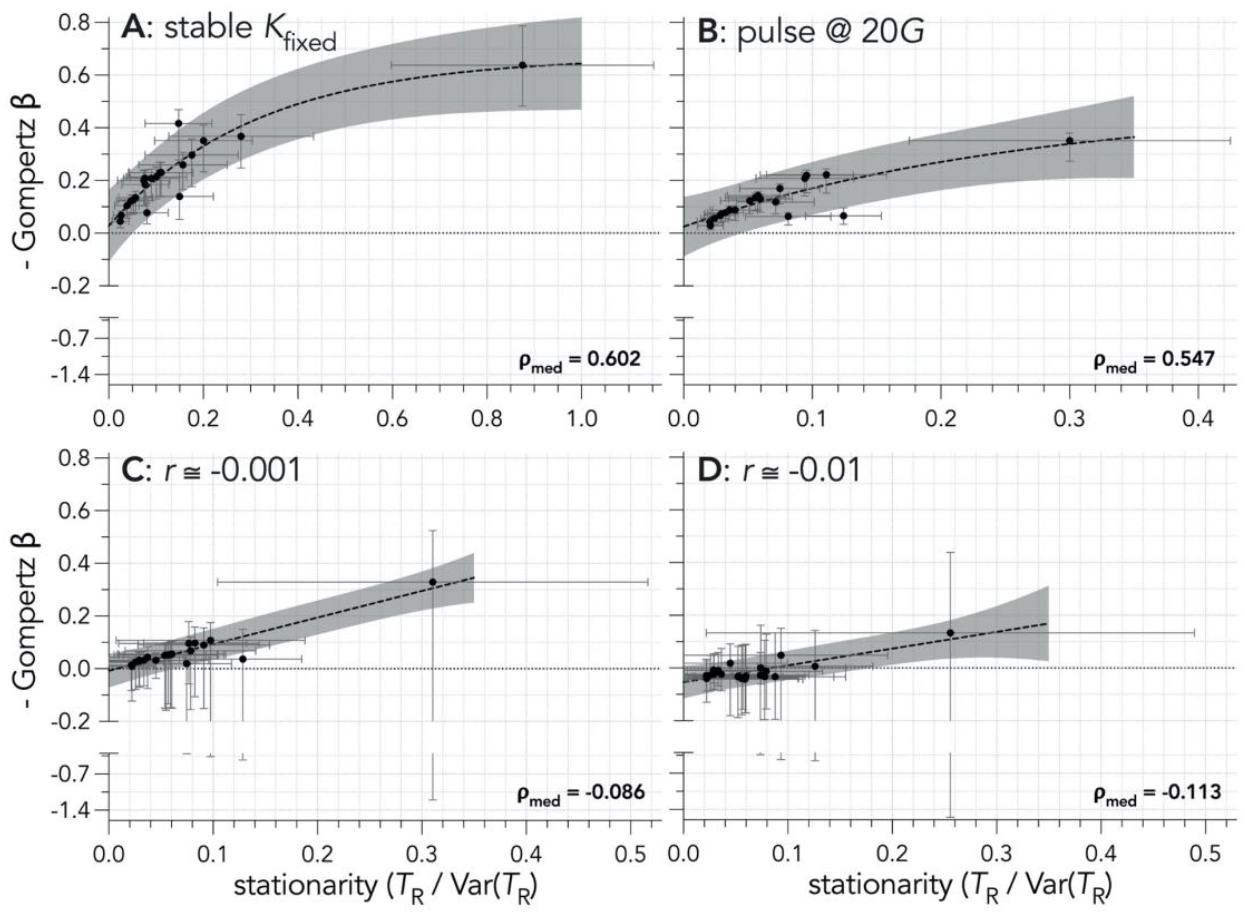
Relationships between the stationarity index 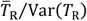 and the strength of ensemble density feedback (slope coefficient *β* × [−1] of the Gompertz-logistic model) for four scenarios with 50% catastrophic (density-indepent) mortality across 21 test species (see Table 1) over 40 generations, including (**A**) carrying capacity (*K*) fixed (scenario 1.2*ii*), (**B**) a pulse disturbance of 90% mortality at 20 generations (20*G*; scenario 1.2*iii*), (**C**) weakly declining (*r* ≅ −0.001, scenario 1.2*iv*), and (**D**) strongly declining (*r* ≅ 0.01, scenario 1.2*v*) populations (scenarios detailed in Table 2). The fitted curves across species exponential plateau models of the form *y* = *y*_max_ - (*y*_max_ - *y*_0_)e*^-kx^* Shaded regions represent the 95% prediction intervals for each type. *ρ*_med_ are the median Spearman’s *ρ* correlation coefficients for the relationship between the ensemble strength and stationarity index across species (resampled 10,000 times; see Fig. S4 for full uncertainty range of *ρ* in each scenario).

**FIGURE 4.**
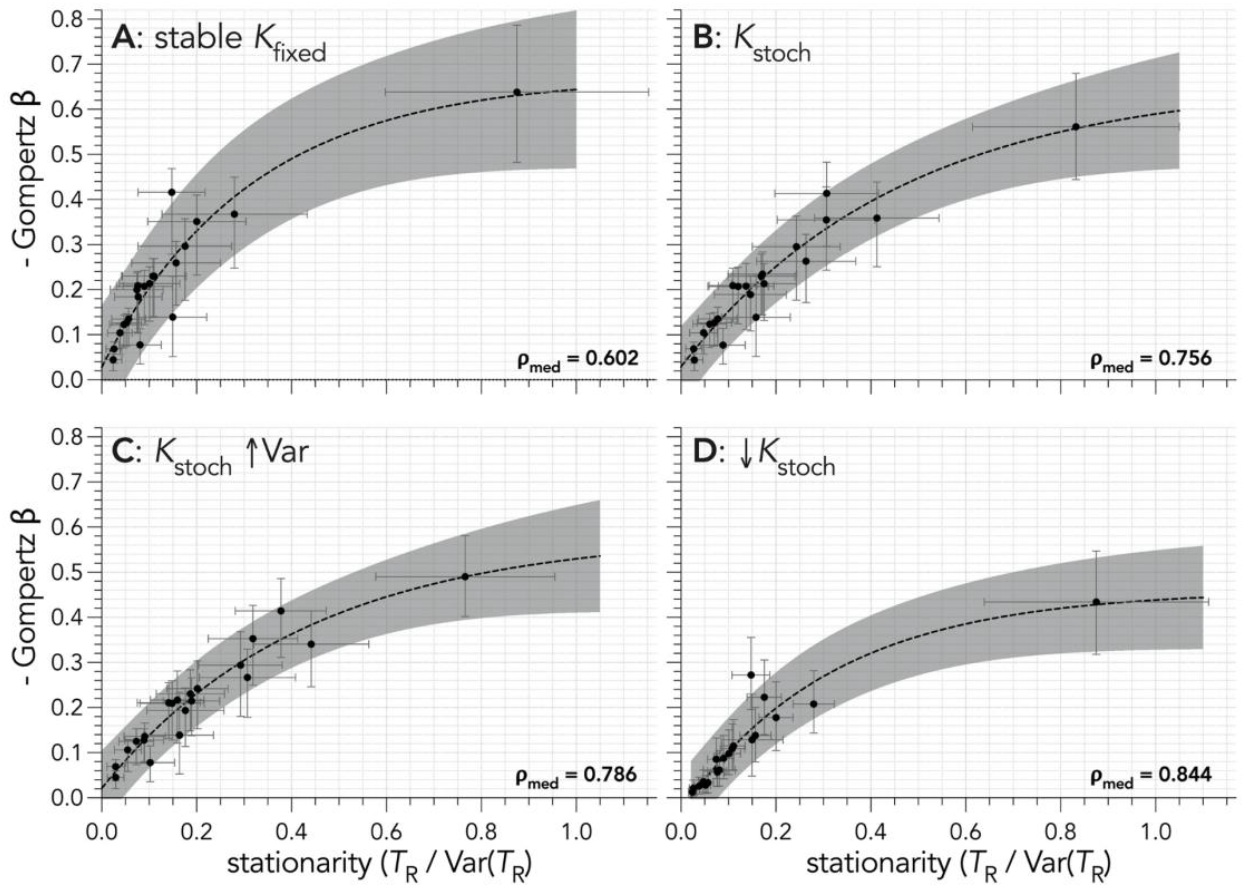
Relationships between the stationarity index 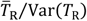 and the strength of ensemble density feedback (slope coefficient *β* × [−1] of the Gompertz-logistic model) across 21 test species (see list in Table 1) over 40 generations for four scenarios (scenarios detailed in Table 2) with 50% catastrophic (density-independent) mortality, including (**A**) carrying capacity (*K*) fixed (scenario 1.2*ii*), (**B**) *K* varying stochastically (*K*_stoch_) around a constant mean with a constant variance (scenario 1.3*vi*), (**C**) *K* varying stochastically with a constant mean and increasing variance (*K*_stoch_ ↑Var, scenario 1.3*vii*), and (**D**) *K* varying stochastically with a declining mean and a constant variance (↓*K*_stoch_, scenario 1.3*viii*). The fitted curves across species exponential plateau models of the form *y* = *y*_max_ - (*y*_max_ - *y*_0_)e*^-kx^* Shaded regions represent the 95% prediction intervals for each type. *ρ*_med_ are the median Spearman’s *ρ* correlation coefficients for the relationship between the ensemble strength and stationarity index across species (resampled 10,000 times; see Fig. S4 for full uncertainty range under each scenario).

## DISCUSSION

Our simulations reveal several new insights into how ensemble (population growth rates) and component (vital rates) density feedbacks can be decoupled. First, the statistical detection of true ensemble feedback strength through phenomenological models is little affected by nonstationarity *per se.* Second, the estimation of ensemble feedback strength through phenomenological models (logistic growth curves; see Introduction) are particularly sensitive to density-independent mortality leading to population decline, but they are less sensitive to moderate fluctuations in carrying capacity. Third, the concern that density-independent processes can invoke false evidence of ensemble signals of compensation are not borne out by our simulations, at least with respect to density-independent mortality.

The mechanisms underlying those trends are nuanced by species’ life histories. For instance, in long-living terrestrial vertebrates (our focus), density feedbacks might operate on fertility to compensate for pathogen-induced adult mortality^50^, those feedbacks might be stronger on survival *versus* fertility when populations are near or far from carrying capacity, respectively^51^, and survival can be entirely driven by climatic conditions and density-independent predation ^52^. In one of the best-studied systems in this regard, Soay sheep from St. Kilda Archipelago (United Kingdom) demonstrate that the demographic role of density and weather varies across sexes and age classes in mild winters, but survival is reduced consistently in all individuals in years of bad weather and high population abundance^53^. Much less-studied than herbivores, inter-pack aggression in carnivores with strong social hierarchies like wolves might shape survival at high densities, but be demographically irrelevant at low densities resulting from prey shortages and/or hunting or culling^54^. Our study lends credence to the application of phenomenological models to the former types of studies addressing the long-term effect of vital rates on population abundance, provided there is enough information available for describing population trends.

Our approach and results do not, of course, explain all possible scenarios leading to the decoupling of ensemble and component feedback signals. For example, many other density-independent factors that we did not consider can dampen the demographic role of social and trophic interactions mediated by population size^2^. Along with the confounding effects of sampling error^55,56^, some of those factors include immigration^57^, spatial heterogeneity in population growth rates^58,59^, fluctuating age structure^60^, and environmental state shifts^29,61,62^. Furthermore, our choice to limit the component mechanisms to feedback on a single demographic rate (albeit, applied to all age classes) for the sake of simpler interpretation could limit the application of our conclusions. For example, additional density-feedback mechanisms operating independently on other demographic rates, such as fertility and dispersal, could potentially complicate the interpretation derived from phenomenological models.

Simulating closed populations potentially inflated the phenomenological model’s capacity to detect the component signal, because permanent dispersal could alleviate per capita reductions in fitness as a population approaches carrying capacity. We also limited our projections to a standardised 40 generations, but even expanding these to 120 generations resulted in little change in the stationarity metric (Fig. S5). Complementary studies focussing on the faster end of the life-history continuum could provide further insights, even though our range of test species still precipitated a life-history signal in terms of component (Fig. S6) and ensemble density-feedback strengths and stationarity (Fig. S7, S8) declining with increasing generation length. However, this relationship faded when the trajectories simulated declines through proportional removal. Indeed, both evidence for^63^ and strength^34^ of ensemble density feedback generally increase along the continuum of slow to fast life histories, because species with slow life histories are assumed to be more demographically stable when density compensation is operating^64^.

While quantifying the true extent of all component density feedback mechanisms operating in real populations will remain challenging in most circumstances, phenomenological models can normally capture the evidence for and strength of the component density feedback mechanism at play. Appreciating the degree of nonstationarity and other types of perturbations affecting abundance time series can contextualise interpretations of ensemble density-feedback signals, especially where substantial density-independent mortality leads to long-term population declines. Importantly, failing to capture density feedback in applied ecological models can lead to suboptimal conservation and management recommendations and outcomes^2,65^.

## Supporting information

Supplementary Results

## ACKNOWLEDGEMENTS

This study was supported by the Australian Research Council through a Centre of Excellence grant (CE170100015) to C.J.A.B. S.H.P. also funded by European Union’s LIFE18 NAT/ES/000121 LIFE DIVAQUA. We acknowledge the Indigenous Traditional Owners of the land on which Flinders University is built — the Kaurna people of the Adelaide Plains.

## AUTHOR CONTRIBUTIONS

CJAB conceived the idea, ran the simulations, and wrote the first draft. SHP reviewed the literature. Both authors contributed to revisions.

## DATA AVAILABILITY STATEMENT

All data files and R code are openly available at https://github/cjabradshaw/DensityFeedbackSims.

## REFERENCES

1 Herrando-Pérez, S., Delean, S., Brook, B. W. & Bradshaw, C. J. A. Density dependence: an ecological Tower of Babel. Oecologia 170, 585–603, doi:10.1007/s00442-012-2347-3 (2012).

2 Herrando-Pérez, S., Delean, S., Brook, B. W. & Bradshaw, C. J. A. Decoupling of component and ensemble density feedbacks in birds and mammals. Ecology 93, 1728–1740, doi:10.1890/11-1415.1 (2012).

3 Eberhardt, L. L., Breiwick, J. M. & Demaster, D. P. Analyzing population growth curves. Oikos 117, 1240–1246 (2008).

4 Fowler, C. W. Density dependence as related to life history strategy. Ecology 62, 602–610 (1981).

5 Matthysen, E. Density-dependent dispersal in birds and mammals. Ecography 28, 403–416 (2005).

6 Brook, B. W. & Bradshaw, C. J. A. Strength of evidence for density dependence in abundance time series of 1198 species. Ecology 87, 1445–1451, doi:10.1890/0012-9658(2006)87%5B1445:SOEFDD%5D2.0.CO;2 (2006).

7 Owen-Smith, N. & Mason, D. R. Comparative changes in adult vs. juvenile survival affecting population trends of African ungulates. J. Anim. Ecol. 74, 762–773, doi:10.1111/j.1365-2656.2005.00973.x (2005).

8 Eberhardt, L. L. A paradigm for population analysis of long-lived vertebrates. Ecology 83, 281–2854 (2002).

9 Paradis, E., Baillie, S. R., Sutherland, W. J. & Gregory, R. D. Exploring density-dependent relationships in demographic parameters in populations of birds at a large spatial scale. Oikos 97, 293–307, doi:10.1034/j.1600-0706.2002.970215.x (2002).

10 Saunders, S. P., Cuthbert, F. J. & Zipkin, E. F. Evaluating population viability and efficacy of conservation management using integrated population models. J. Appl. Ecol. 55, 1380–1392, doi:10.1111/1365-2664.13080 (2018).

11 Doyle, S. et al. Temperature and precipitation at migratory grounds influence demographic trends of an Arctic-breeding bird. Glob. Change Biol. 26, 5447–5458, doi:10.1111/gcb.15267 (2020).

12 Margalida, A. et al. An assessment of population size and demographic drivers of the Bearded Vulture using integrated population models. Ecol Monogr 90, e01414, doi:10.1002/ecm.1414 (2020).

13 Morrison, C. A. et al. Covariation in population trends and demography reveals targets for conservation action. Proc. R. Soc. Lond. B 288, 20202955, doi:10.1098/rspb.2020.2955 (2021).

14 Pardo, D. et al. Additive effects of climate and fisheries drive ongoing declines in multiple albatross species. Proc. Natl. Acad. Sci. U.S.A. 114, E10829–E10837, doi:10.1073/pnas.1618819114 (2017).

15 Stillman, R. A. et al. Predicting impacts of food competition, climate, and disturbance on a long-distance migratory herbivore. Ecosphere 12, doi:10.1002/ecs2.3405 (2021).

16 Azerefegne, F., Solbreck, C. & Ives, A. R. Environmental forcing and high amplitude fluctuations in the population dynamics of the tropical butterfly *Acraea acerata* (Lepidoptera: Nymphalidae). J. Anim. Ecol. 70, 1032–1045, doi:10.1046/j.0021-8790.2001.00556.x (2001).

17 Jepsen, J. U., Hagen, S. B., Karlsen, S. R. & Ims, R. A. Phase-dependent outbreak dynamics of geometrid moth linked to host plant phenology. Proc. R. Soc. Lond. B 276, 4119–4128, doi:10.1098/rspb.2009.1148 (2009).

18 Bonsall, M. B. & Benmayor, R. Multiple infections alter density dependence in host-pathogen interactions. J. Anim. Ecol. 74, 937–945, doi:10.1111/j.1365-2656.2005.00991.x (2005).

19 Ma, Z. A unified survival-analysis approach to insect population development and survival times. Sci. Rep. 11, 8223, doi:10.1038/s41598-021-87264-1 (2021).

20 Marini, G. et al. The role of climatic and density dependent factors in shaping mosquito population dynamics: the case of *Culex pipiens* in northwestern Italy. PLoS One 11, e0154018, doi:10.1371/journal.pone.0154018 (2016).

21 Maud, J. L. et al. How does *Calanus helgolandicus* maintain its population in a variable environment? Analysis of a 25-year time series from the English Channel. Progress in Oceanography 137, 513–523, doi:10.1016/j.pocean.2015.04.028 (2015).

22 McGeoch, M. A. & Price, P. W. Scale-dependent mechanisms in the population dynamics of an insect herbivore. Oecologia 144, 278–288, doi:10.1007/s00442-005-0073-9 (2005).

23 Bellier, E., Kéry, M. & Schaub, M. Simulation-based assessment of dynamic N-mixture models in the presence of density dependence and environmental stochasticity. Meth Ecol Evol 7, 1029–1040, doi:10.1111/2041-210X.12572 (2016).

24 Berryman, A. & Turchin, P. Identifying the density-dependent structure underlying ecological time series. Oikos 92, 265–270 (2001).

25 Münster-Swendsen, M. & Berryman, A. Detecting the causes of population cycles by analysis of R-functions: the spruce needle-miner, *Epinotia tedella*, and its parasitoids in Danish spruce plantations. Oikos 108, 495–502, doi:10.1111/j.0030-1299.2005.13747.x (2005).

26 Kolb, A., Dahlgren, J. P. & Ehrlén, J. Population size affects vital rates but not population growth rate of a perennial plant. Ecology 91, 3210–3217, doi:10.1890/09-2207.1 (2010).

27 Bürgi, L. P., Roltsch, W. J. & Mills, N. J. Allee effects and population regulation: a test for biotic resistance against an invasive leafroller by resident parasitoids. Popul. Ecol. 57, 215–225, doi:10.1007/s10144-014-0451-4 (2015).

28 Battaile, B. C. & Trites, A. W. Linking reproduction and survival can improve model estimates of vital rates derived from limited time-series counts of pinnipeds and other species. PLoS One 8, e77389, doi:10.1371/journal.pone.0077389 (2013).

29 Turchin, P. Complex Population Dynamics: A Theoretical/Empirical Synthesis. (Princeton University Press, 2003).

30 Heppell, S. S., Caswell, H. & Crowder, L. B. Life histories and elasticity patterns: perturbation analysis for species with minimal demographic data. Ecology 81, 654–665 (2000).

31 Oli, M. K. & Dobson, F. S. The relative importance of life-history varibales to population growth rate in mammals: Cole’s predictions revisited. Am. Nat. 161, 422–440 (2003).

32 Sæther, B.-E. & Bakke, Ø. Avian life history variation and contribution of demographic traits to the population growth rate. Ecology 81, 642–653, doi:10.1890/0012-9658(2000)081[0642:ALHVAC]2.0.CO;2 (2000).

33 Bradshaw, C. J. A. et al. Relative demographic susceptibility does not explain the extinction chronology of Sahul’s megafauna. eLife 10, e63870, doi:10.7554/eLife.63870 (2021).

34 Herrando-Pérez, S., Delean, S., Brook, B. W. & Bradshaw, C. J. A. Strength of density feedback in census data increases from slow to fast life histories. Ecol Evol 2, 1922–1934, doi:10.1002/ece3.298 (2012).

35 Gaillard, J. M. et al. An analysis of demographic tactics in birds and mammals. Oikos 56, 59–76 (1989).

36 Oakwood, M., Bradley, A. J. & Cockburn, A. Semelparity in a large marsupial. Proc. R. Soc. Lond. B 268, 407–411 (2001).

37 Cockburn, A. in Marsupial Biology: Recent Research, New Perspectives (eds N. Saunders & L. Hinds) 163–171 (University of New South Wales Press, 1997).

38 Holz, P. H. & Little, P. B. Degenerative leukoencephalopathy and myelopathy in dasyurids. J. Wildl. Dis. 31, 509–513 (1995).

39 Caswell, H. Matrix Population Models: Construction, Analysis, and Interpretation, 2nd edn. (Sinauer Associates, Inc., 2001).

40 Traill, L. W., Brook, B. W., Frankham, R. & Bradshaw, C. J. A. Pragmatic population viability targets in a rapidly changing world. Biol. Conserv. 143, 28–34, doi:10.1016/j.biocon.2009.09.001 (2010).

41 Brook, B. W., Traill, L. W. & Bradshaw, C. J. A. Minimum viable population size and global extinction risk are unrelated. Ecol. Lett. 9, 375–382 (2006).

42 Reed, D. H., O’Grady, J. J., Ballou, J. D. & Frankham, R. The frequency and severity of catastrophic die-offs in vertebrates. Anim. Conserv. 6, 109–114, doi:10.1017/S1367943003147 (2003).

43 Bradshaw, C. J. A. et al. More analytical bite in estimating targets for shark harvest. Mar. Ecol. Prog. Ser. 488, 221–232, doi:10.3354/meps10375 (2013).

44 Berryman, A. A. Principles of Population Dynamics and Their Application. (Stanley Thorners Ltd., 1999).

45 Verhulst, P. F. Notice sur la loi que la population poursuit dans son accroissement. Correspondance mathématique et physique 10, 113–121 (1838).

46 Ricker, W. E. Stock and recruitment. J Fish Res Board Can 11, 559–623 (1954).

47 Nelder, J. A. The fitting of a generalization of the logistic curve. Biometrics 17, 89–110 (1961).

48 Burnham, K. P. & Anderson, D. R. Model Selection and Multimodel Inference: A Practical Information-Theoretic Approach. 2nd edn, (Springer-Verlag, 2002).

49 Doncaster, C. P. Non-linear density dependence in time series is not evidence of non-logistic growth. Theor Popul Biol 73, 483–489 (2008).

50 McDonald, J. L. et al. Demographic buffering and compensatory recruitment promotes the persistence of disease in a wildlife population. Ecol. Lett. 19, 443–449, doi:10.1111/ele.12578 (2016).

51 Sæther, B. E. et al. Demographic routes to variability and regulation in bird populations. Nat. Comm. 7, 12001, doi:10.1038/ncomms12001 (2016).

52 Hebblewhite, M., Eacker, D. R., Eggeman, S., Bohm, H. & Merrill, E. H. Density-independent predation affects migrants and residents equally in a declining partially migratory elk population. Oikos 127, 1304–1318, doi:10.1111/oik.05304 (2018).

53 Coulson, T. et al. Age, sex, density, winter weather and population crashes in soay sheep. Science 292, 1528–1531 (2001).

54 Cubaynes, S. et al. Density-dependent intraspecific aggression regulates survival in northern Yellowstone wolves *(Canis lupus)*. J. Anim. Ecol. 83, 1344–1356, doi:10.1111/1365-2656.12238 (2014).

55 Staples, D. F., Taper, M. L. & Dennis, B. Estimating population trend and process variation for PVA in the presence of sampling error. Ecology 85, 923–929, doi:10.1890/03-3101 (2004).

56 Knape, J. & de Valpine, P. Are patterns of density dependence in the Global Population Dynamics Database driven by uncertainty about population abundance? Ecol. Lett. 15, 17–23, doi:10.1111/j.1461-0248.2011.01702.x (2012).

57 Lieury, N. et al. Compensatory immigration challenges predator control: an experimental evidence-based approach improves management. J Wildl Manag 79, 425–434, doi:10.1002/jwmg.850 (2015).

58 Thorson, J. T. et al. The importance of spatial models for estimating the strength of density dependence. Ecology 96, 1202–1212, doi:10.1890/14-0739.1 (2015).

59 Johnson, D. W., Freiwald, J. & Bernardi, G. Genetic diversity affects the strength of population regulation in a marine fish. Ecology 97, 627–639, doi:10.1890/15-0914 (2016).

60 Hoy, S. R. et al. Fluctuations in age structure and their variable influence on population growth. Funct. Ecol. 34, 203–216, doi:10.1111/1365-2435.13431 (2020).

61 Lande, R. et al. Estimating density dependence from population time series using demographic theory and life-history data. Am. Nat. 159, 321–337 (2002).

62 Wu, Z., Huang, N. E., Long, S. R. & Peng, C.-K. On the trend, detrending, and variability of nonlinear and nonstationary time series. Proc. Natl. Acad. Sci. U.S.A. 104, 14889, doi:10.1073/pnas.0701020104 (2007).

63 Holyoak, M. & Baillie, S. R. Factors influencing detection of density dependence in British birds. II. Longevity and population variability. Oecologia 108, 54–63 (1996).

64 Sæther, B.-E., Engen, S. & Matthysen, E. Demographic characteristics and population dynamical patterns of solitary birds. Science 295, 2070–2073 (2002).

65 Horswill, C., O’Brien, S. H. & Robinson, R. A. Density dependence and marine bird populations: are wind farm assessments precautionary? J. Appl. Ecol. 54, 1406–1414, doi:10.1111/1365-2664.12841 (2017).

