## Supplementary Results for "Density-independent processes decouple component and ensemble density feedbacks"

### SUPPLEMENTARY INFORMATION

**FIGURE S1** Probability of an ensemble compensatory density-feedback signal (Pr(DF) = $\Sigma w\text{AIC}_c$-DF = sum of Akaike's information criterion weights across the Ricker- and Gompertz-logistic models — see Materials and methods) in abundance time series for simulated populations of 21 long-lived species of Australian mammals and birds (see list in Table 1) subjected to compensatory density feedback on survival and experiencing 50 % catastrophic (density-independent) mortality over 40 generations. Each probability surface represents one of the 21 test species (see list in Table 1), so plots show the overlapping median probability density over 10,000 times series of abundance per species and for each of four demographic scenarios (detailed in Table 2), including (**A**) a carrying capacity is fixed ($K_{\text{fixed}}$) with 50 % catastrophic (density-independent) mortality (scenario 1.2*ii*), (**B**) a pulse disturbance of 90% mortality at 20 generations (20*G*; scenario 1.2*iii*), and (**C**) weakly declining ($\bar{r}$ ≅ -0.001; scenario 1.2*iv*) and (D) strongly declining ($\bar{r}$ ≅ 0.01; scenario 1.2*v*) populations. See Fig. S3 for bootstrapped mean Spearman correlation coefficients for each scenario.

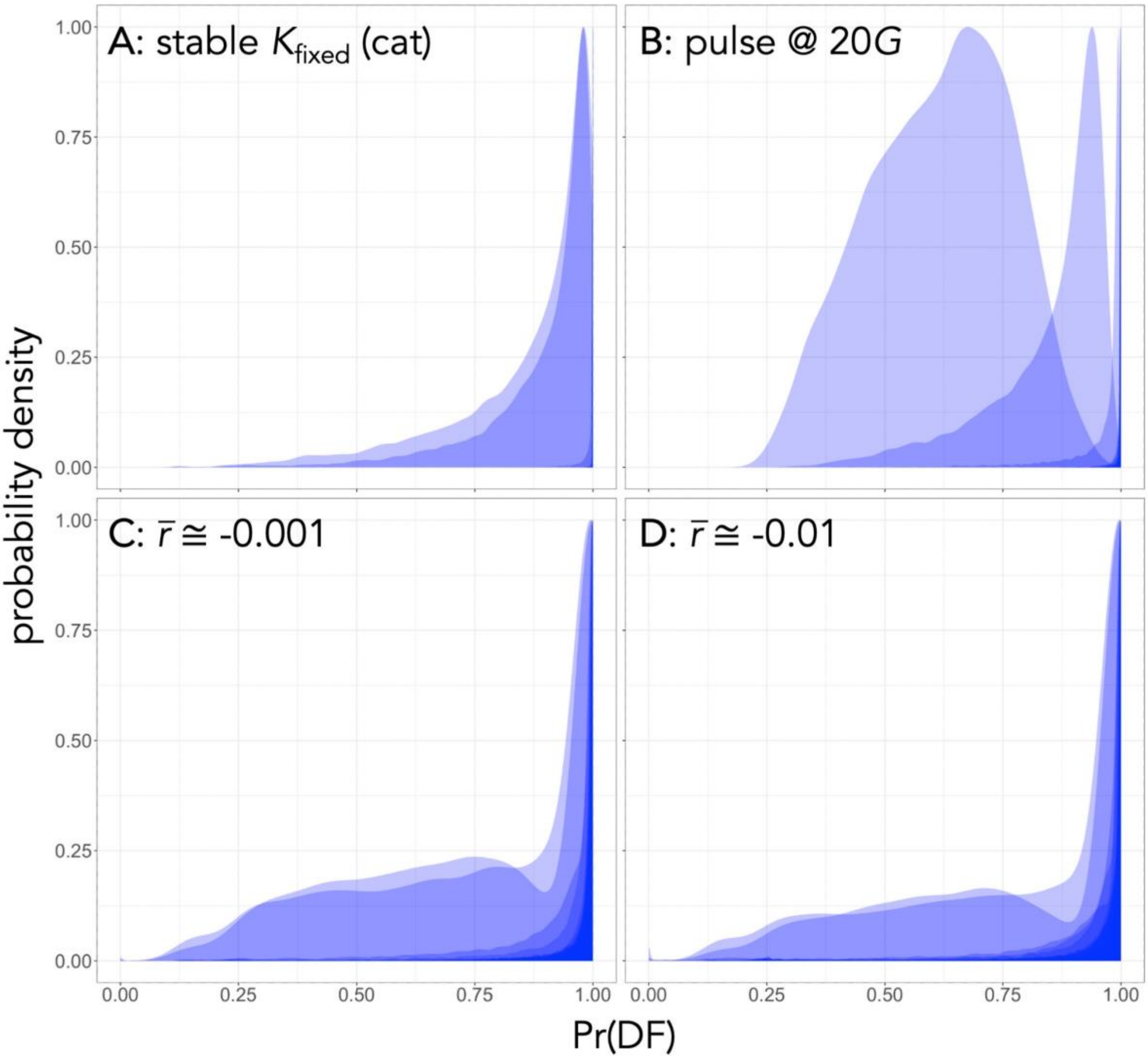

**FIGURE S2** Probability of an ensemble compensatory density-feedback signal (Pr(DF) = Σ*w*AIC$_c$-DF = sum of Akaike's information criterion weights across the Ricker- and Gompertz-logistic models — see Materials and methods) in abundance time series for simulated populations of 21 long-lived species of Australian mammals and birds (see list in Table 1) subjected to compensatory density feedback on survival and experiencing fluctuations in carrying capacity (*K*) along with 50 % catastrophic (density-independent) mortality over 40 generations. Each probability surface represents one of the 21 test species (see list in Table 1), so plots show the overlapping median probability density over 10,000 times series of abundance per species and for each of four demographic scenarios (detailed in Table 2), including (**A**) a stable demographic projection where *K* is fixed ($K_{\text{fixed}}$) (scenario 1.2*ii*), (B) *K* varies stochastically ($K_{\text{stoch}}$) around a constant mean with a constant variance (scenario 1.3*vi*), (**C**) *K* varying stochastically with a constant mean and increasing variance ($K_{\text{stoch}}$↑Var; scenario 1.3*vii*), and (**C**) *K* varying stochastically with a declining mean and a constant variance (↓$K_{\text{stoch}}$; scenario 1.3*viii*). See Fig. S3 for bootstrapped mean Spearman correlation coefficients for each scenario.

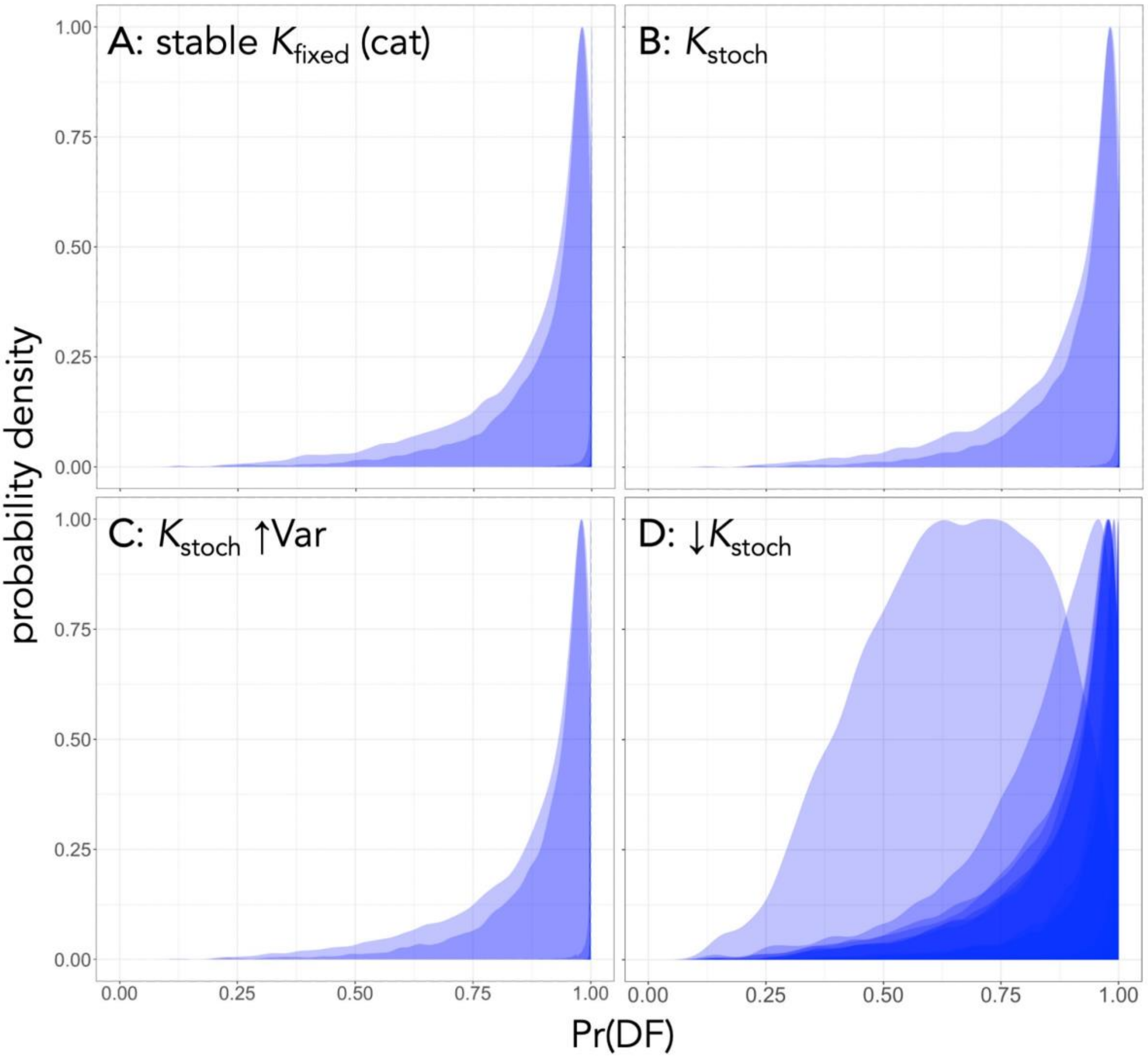

**FIGURE S3** Bootstrapped mean (with 80 % confidence intervals; 100,000 resamples) probability of an ensemble compensatory density-feedback signal (Pr(DF) = $\Sigma w\text{AIC}_c$-DF = sum of Akaike's information criterion weights across the Ricker- and Gompertz-logistic models — see Materials and methods) in abundance time series for simulated populations of 21 long-lived species of Australian mammals and birds for populations (see list in Table 1) subjected to compensatory density feedback on survival and experiencing fluctuations in carrying capacity (*K*) and/or 50 % catastrophic (density-independent) mortality (scenarios detailed in Table 2). Demographic scenarios (see details in Table 2) include (**A**) *K* fixed ($K_{\text{fixed}}$) with no catastrophic mortality (no cat; scenario 1.1*i*), and with catastrophic mortality in combination with (**B**) $K_{\text{fixed}}$ (cat; scenario 1.2*ii*), (**C**) a pulse disturbance of 90% mortality at 20 generations (20*G*; scenario 1.2*iii*), (**D**) weakly declining ($\bar{r} \cong -0.001$; scenario 1.2*iv*) and (**E**) strongly declining ($\bar{r} \cong 0.01$; scenario 1.2*v*) populations, (**F**) *K* varying stochastically($K_{\text{stoch}}$) around a constant mean with a constant variance (scenario 1.3*vi*), (**G**) *K* varying stochastically with a constant mean and increasing variance ($K_{\text{stoch}}\uparrow$Var; scenario 1.3*vii*), and (**H**) *K* varying stochastically with a declining mean and a constant variance ($\downarrow K_{\text{stoch}}$; scenario 1.3*viii*). The vertical dashed line at Pr(DF) = 0.5 in each panel is the point below which the evidence for a density-independent model [Pr(DI) = $\Sigma w\text{AIC}_c$-DI = sum of Akaike's information criterion weights across the random walk and exponential models] is greater than Pr(DF). See Table 2 for species abbreviations.

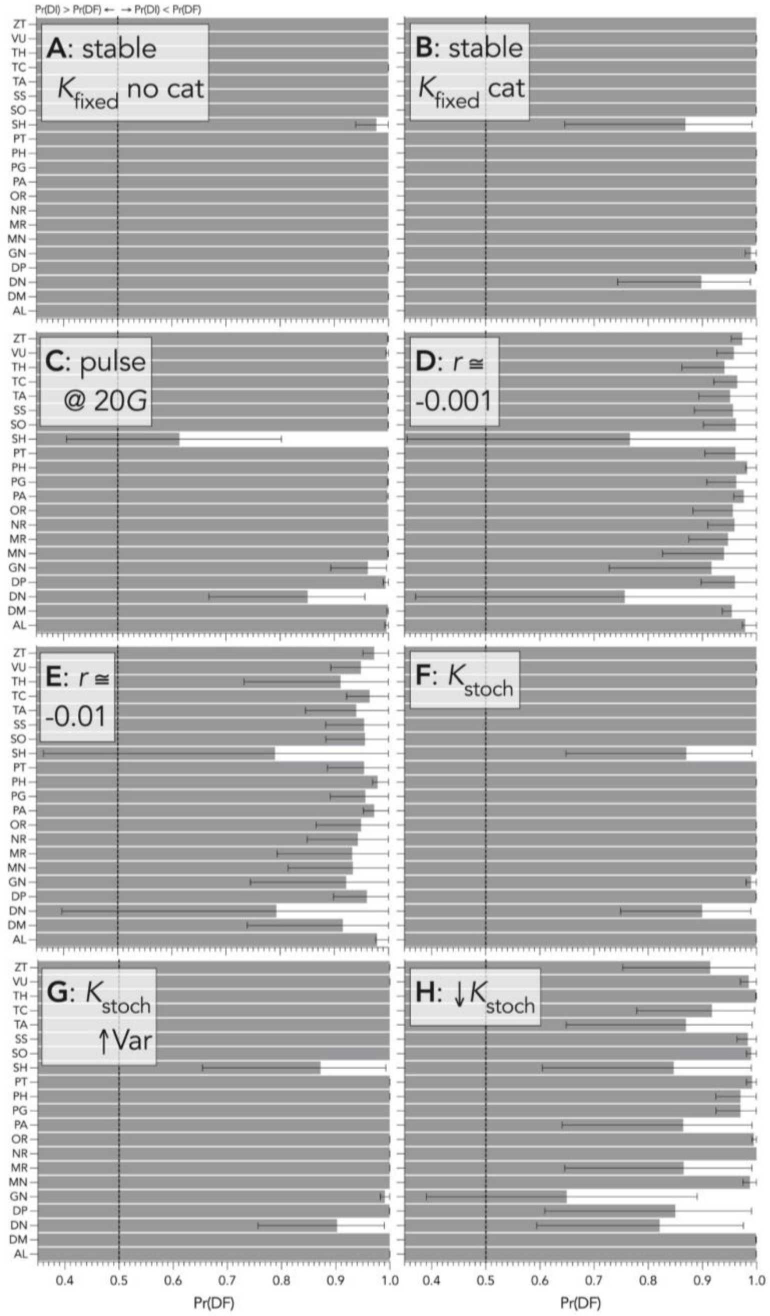

**FIGURE S4** Bootstrapped (10,000 iterations) Spearman's correlation $\rho$ between (**A**) ensemble density feedback strength (- Gompertz slope $\beta$, the reduction of survival as population density increases) *and* component feedback strength on survival ($1 - S_{red}$, the reduction in survival as population density increases), and (**B**) ensemble feedback strength *and* the stationarity metric $\bar{T}_R/\mathrm{Var}(T_R)$ for 10,000 simulated populations across each of 21 long-lived species of Australian mammals and birds for populations (see list in Table 1) subjected to compensatory density feedback on survival and experiencing fluctuations in carrying capacity ($K$) and/or 50 % catastrophic (density-independent) mortality (scenarios detailed in Table 2) . Demographic scenarios include $K$ fixed ($K_{fixed}$) with no catastrophic mortality (no cat; scenario 1.1*i*), and catastrophic mortality in combination with $K_{fixed}$ (cat; scenario 1.2*ii*), a pulse disturbance of 90% mortality at 20 generations (20*G*; scenario 1.2*iii*), weakly declining ($\bar{r} \cong$ -0.001; scenario 1.2*iv*) and (**E**) strongly declining ($\bar{r} \cong$ 0.01; scenario 1.2*v*) populations, $K$ varying stochastically($K_{stoch}$) around a constant mean with a constant variance (scenario 1.3*vi*), $K$ varying stochastically with a constant mean and increasing variance ($K_{stoch}\uparrow$Var; scenario 1.3*vii*), and $K$ varying stochastically with a declining mean and a constant variance ($\downarrow K_{stoch}$; scenario 1.3*viii*).

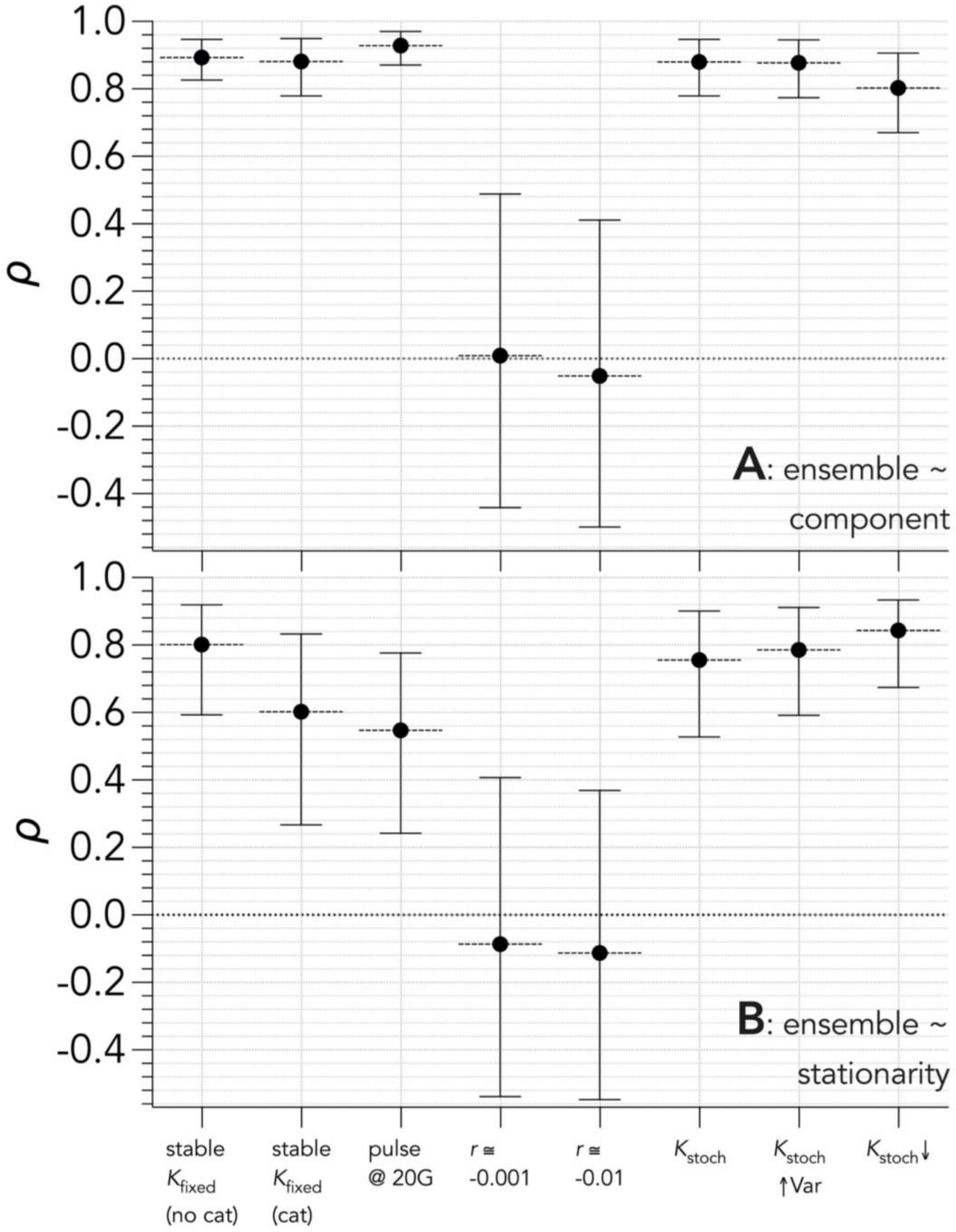

**FIGURE S5** Truncated violin plots showing the distribution of the stationarity index $\bar{T}_{\mathrm{R}}/\mathrm{Var}(T_{\mathrm{R}})$ across 10,000 time series of population abundance per species and all 21 species (see species list in Table 1) obtained from age-structured populations for scenarios showing carrying capacity fixed with component compensatory density-feedback on survival and 50% catastrophic (density-independent) mortality to produce stable population growth rates around 0 over 40 (scenario 1.2*ii*; detailed in Table 2) and 120 generations (*G*).

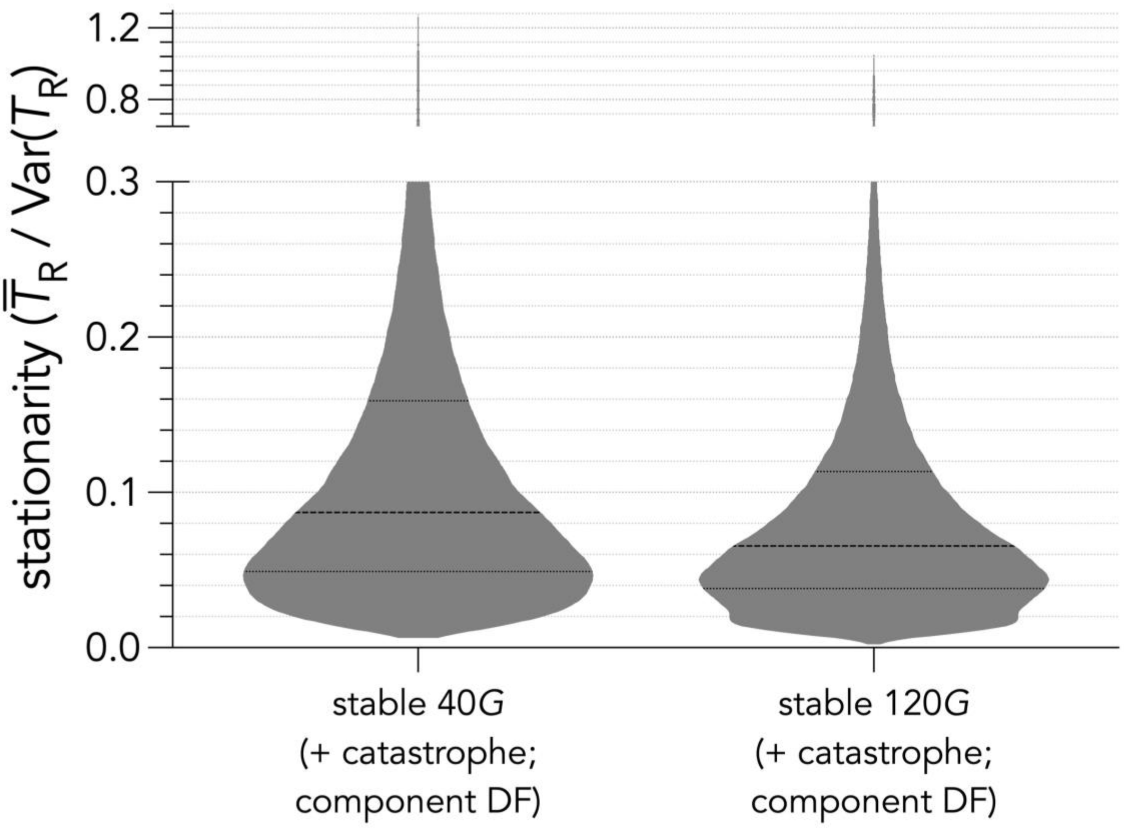

**Fig. S6**. Relationship between strength of component density feedback and generation length (years) across 10,000 time series of population abundance for each of 21 test species (see list in Table 1) obtained from age-structured populations subjected to a compensatory component density feedback on survival over 40 generations for a demographic scenario with constant carrying capacity and no catastrophic (density-independent) mortality (scenario 1.1*i*; detailed in Table 2). The dashed grey line indicates a least-squares-fitted (adjusted coefficient of regression $R^2$ = 0.58) exponential plateau model of the form: $y = y_{max} - (y_{max} - y_0)e^{-kG}$, where $y_0$ = starting value of component strength, $y_{max}$ = maximum component strength, $k$ = rate constant (years$^{-1}$) and $G$ = generation time (years). Species notation: DP = *Diprotodon optatum*, PA = *Palorchestes azael*, ZT = *Zygomaturus trilobus*, PH = *Phascolonus gigas*, VU *Vombatus ursinus* (herbivore vombatiform); PG = *Procoptodon goliah*, SS = *Sthenurus stirlingi*, PT = *Protemnodon anak*, SO = *Simosthenurus occidentalis*, MN = *Metasthenurus newtonae*, OR = *Osphranter rufus* (herbivore macropodiformes); GN = *Genyornis newtoni*, DN = *Dromaius novaehollandiae* (large omnivore birds) , AL = *Alectura lathami*; TC = *Thylacoleo carnifex*, TH = *Thylacinus cynocephalus*, SH = *Sarcophilus harrisii* (carnivores) , DM = *Dasyurus maculatus*; MR = *Megalibgwilia ramsayi*; TA = *Tachyglossus aculeatus* (invertivore monotremes).

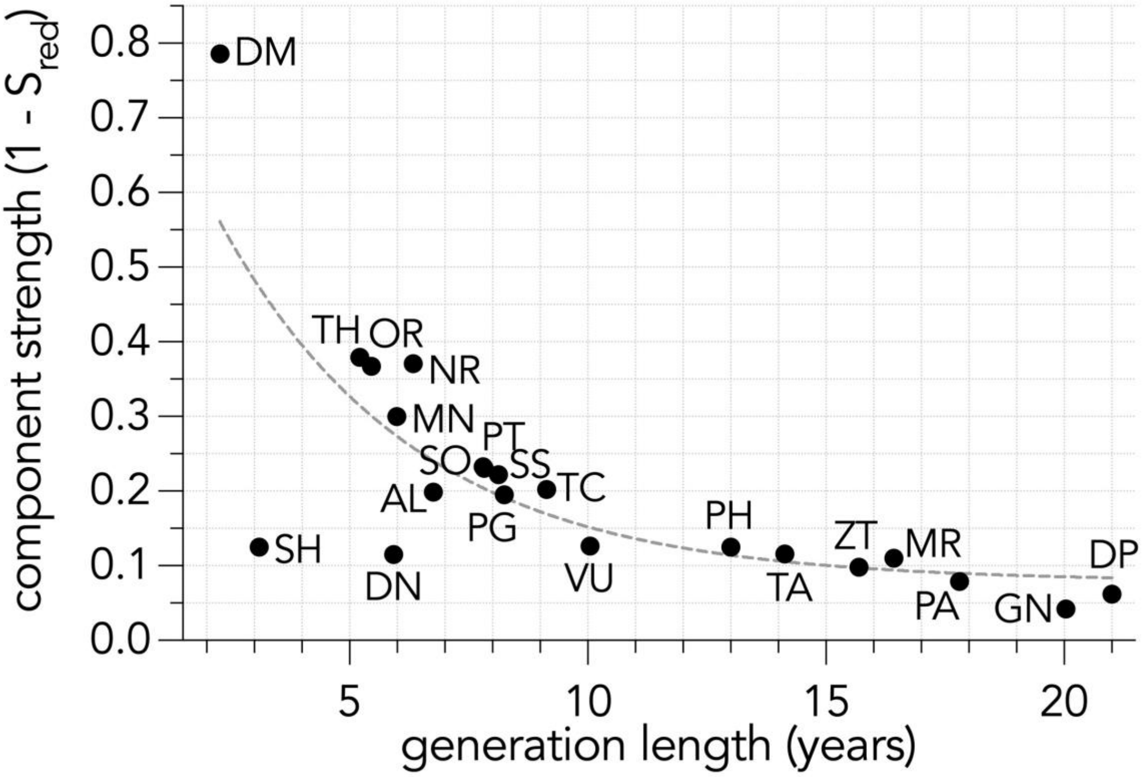

**FIGURE S7** Relationships between the stationarity index $\bar{T}_{\mathrm{R}}/\mathrm{Var}(T_{\mathrm{R}})$ and generation length across 10,000 times series of population abundance per species and all 21 test species (see list in Table 1) obtained from age-structured populations subjected to a compensatory component density feedback on survival over 40 generations, according to seven demographic scenarios (detailed in Table 2). Demographic scenarios include (**A**) carrying capacity *K* fixed ($K_{\mathrm{fixed}}$; scenario 1.2*ii*), (**B**) a pulse disturbance of 90% mortality at 20 generations (20G; scenario 1.2*iii*), (**C**) weakly declining ($\bar{r} \cong$ -0.001; scenario 1.2*iv*) and (**D**) strongly declining ($\bar{r} \cong$ 0.01; scenario 1.2*v*) populations, (**E**) *K* varying stochastically ($K_{\mathrm{stoch}}$) around a constant mean with a constant variance (scenario 1.3*vi*), (**F**) *K* varying stochastically with a constant mean and increasing variance ($K_{\mathrm{stoch}}\uparrow$Var; scenario 1.3*vii*), and (**G**) *K* varying stochastically with a declining mean and a constant variance ($\downarrow K_{\mathrm{stoch}}$; scenario 1.3*viii*).

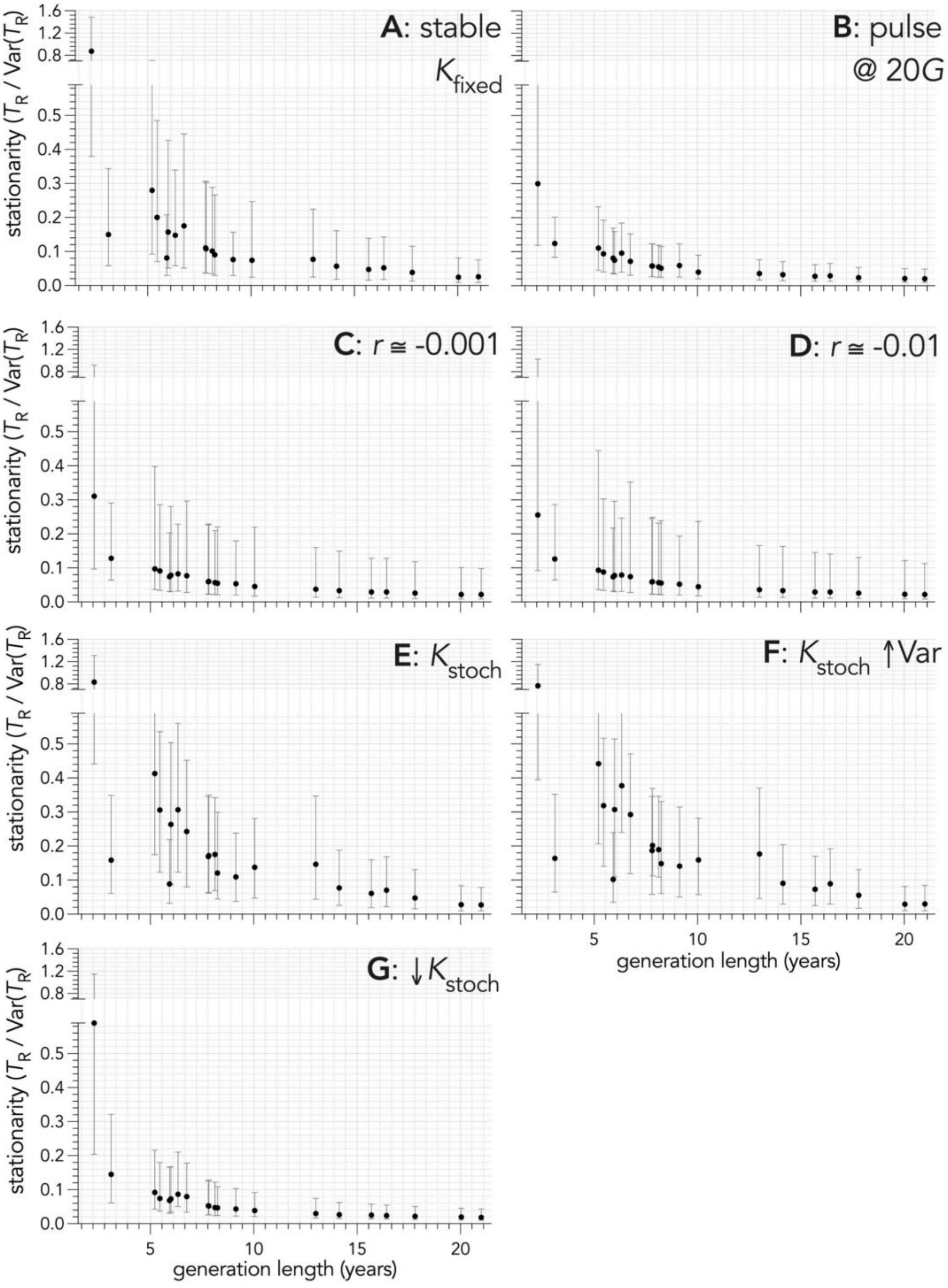

**FIGURE S8** Relationships between the strength of ensemble (- Gompertz slope $\beta$, the reduction of survival as population density increases) and generation length across 10,000 times series of population abundance per species and all 21 test species (see list in Table 1) obtained from age-structured populations subjected to a compensatory component density feedback on survival over 40 generations, according to seven demographic scenarios (detailed in Table 2). Demographic scenarios include (**A**) carrying capacity $K$ fixed ($K_{\text{fixed}}$; scenario 1.2*ii*), (**B**) a pulse disturbance of 90% mortality at 20 generations (20$G$; scenario 1.2*iii*), (**C**) weakly declining ($\bar{r} \cong -0.001$; scenario 1.2*iv*) and (**D**) strongly declining ($\bar{r} \cong 0.01$; scenario 1.2*v*) populations, (**E**) $K$ varying stochastically ($K_{\text{stoch}}$) around a constant mean with a constant variance (scenario 1.3*vi*), (**F**) K varying stochastically with a constant mean and increasing variance ($K_{\text{stoch}}$↑Var; scenario 1.3*vii*), and (**G**) $K$ varying stochastically with a declining mean and a constant variance (↓$K_{\text{stoch}}$; scenario 1.3*viii*).

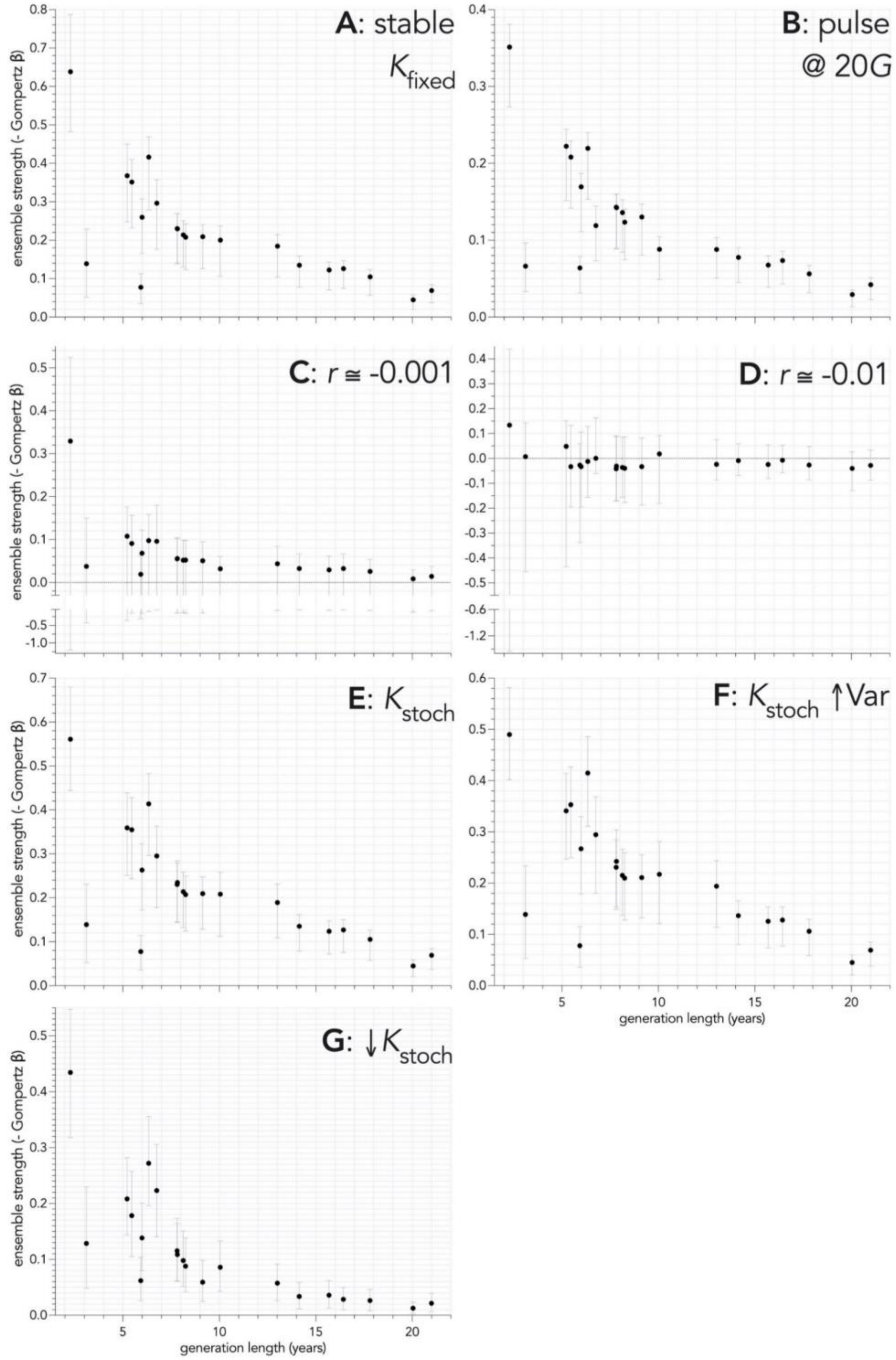
